# CHAF1A promotes the translesion DNA synthesis pathway in response to DNA replication stress

**DOI:** 10.1101/2023.04.21.537900

**Authors:** Bing Wen, Hai-Xiang Zheng, Dan-Xia Deng, Zhi-Da Zhang, Jing-Hua Heng, Lian-Di Liao, Li-Yan Xu, En-Min Li

## Abstract

The translesion DNA synthesis (TLS) pathway mediated by proliferating cell nuclear antigen (PCNA) monoubiquitination is an essential mechanism by which cancer cells bypass DNA damage caused by DNA replication stress to maintain genomic stability and cell survival. Chromatin assembly factor 1 subunit A (CHAF1A) traditionally promotes histone assembly during DNA replication. Here, we revealed that CHAF1A is a novel regulator of the TLS pathway. High expression of CHAF1A is significantly associated with poor prognosis in cancer patients. CHAF1A promotes fork restart under DNA replication stress and maintains genome integrity. CHAF1A enhances the interaction between PCNA and E3 ubiquitin protein ligase RAD18 and promotes PCNA monoubiquitination, thereby promoting the recruitment of Y-family DNA polymerase Pol η and enhancing cancer cell resistance to stimuli that trigger replication fork blockade. Mechanistically, CHAF1A-mediated PCNA monoubiquitination is independent of CHAF1A-PCNA interaction. CHAF1A interacts with both RAD18 and replication protein A2 (RPA2), mediating RAD18 binding on chromatin in response to DNA replication stress. Taken together, these findings improve our understanding of the mechanisms that regulate the TLS pathway and provide insights into the relationship between CHAF1A and the malignant progression of cancers.

## Introduction

The faithful transmission of genetic information to daughter cells is essential for maintaining the genomic integrity in cancer cells and depends on complete and accurate DNA replication during the S-phase of cell division ^1^. However, endogenous and exogenous stimuli can cause DNA lesions that prevent accurate DNA replication, causing DNA replication stress ^2, 3^. Since the correct processing of stalled or damaged replication forks is critical for maintaining genomic stability and ensuring cell survival ^4^, cancer cells have developed mechanisms to resist DNA replication stress and promote DNA repair to combat genomic instability ^5^.

During replication, at the center of the replication fork, three proliferating cell nuclear antigen (PCNA) monomers form a sliding clamp that plays an important role in the progression of DNA replication and provides a central scaffold that allows dynamic interactions between the replication machinery and numerous additional factors ^6, 7^. In mammalian cells, the predominant mechanism used to bypass DNA damage is the translesion synthesis (TLS) pathway, which is mediated by monoubiquitinated PCNA ^8^. DNA replication stress inducers such as hydroxyurea (HU), methyl methanesulfonate (MMS), cisplatin, aphidicolin (APH), and ultraviolet (UV) radiation can dramatically increase PCNA monoubiquitination at Lys164 (K164) by the RAD6/RAD18 complex, resulting in the stalling of DNA replication forks ^9–14^. In addition, PCNA monoubiquitination promotes the conversion of replicative DNA polymerase Pol δ/ε into damage-tolerant translesion (DTT) DNA polymerases, such as Pol η, Pol κ, or REV1 ^7^. TLS pathway activation is therefore beneficial for cancer cell survival under DNA replication stress and is closely related to the malignant progression of cancers and resistance to chemotherapy ^15, 16^.

CAF-1 is a heterotrimeric histone chaperone complex that consists of three subunits: chromatin assembly factor 1 subunit A (CHAF1A, also known as p150 or CAF1A), chromatin assembly factor 1 subunit B (CHAF1B, also known as p60 or CAF1B), and histone-binding protein RBBP4 (also known as p48) ^17^. CAF-1 mainly binds to newly synthesized histones (H3 especially H3.1/H3.2, and H4) by interacting with PCNA and promotes their deposition at the replication fork during S-phase. In addition, CHAF1A recruits heterochromatin protein 1 homolog alpha (HP1α) to DNA damage sites and promotes homologous recombination repair ^18^. Although CHAF1A and PCNA have been shown to co-localize at UV radiation-damaged sites ^19^, their role in DNA damage tolerance pathway remains unclear. Here we demonstrate a novel function of CHAF1A in regulating PCNA monoubiquitination mediated-TLS pathway and contributing to cancer cell genome integrity and cancer progression.

## Materials and methods

### Cell culture

HEK293T cells and A549 cells were purchased from American Type Culture Collection (ATCC, Manassas, VA, USA). We obtained the human esophageal cancer cell lines KYSE30 and KYSE510 that were established by Dr. Shimada Yutaka (Faculty of Medicine, Kyoto University, Kyoto, Japan) ^20^. All cells were cultured in RPMI-1640 medium supplemented with 10% fetal bovine serum (FBS), penicillin (100 mg/mL), and streptomycin (100 mg/mL) in a humidified 5% CO_2_ atmosphere at 37 °C and were tested for mycoplasma contamination using short tandem repeat (STR) analysis (IGEbio, Guangzhou, China).

### Generation of cell lines with inducible CHAF1A knockdown

Cells with inducible CHAF1A knockdown were generated using a lentiviral system ^21^. First, we purchased shCHAF1A oligos (shCHAF1A#1 oligo1: 5ʹ-CCG GCC GAC TCA ATT CCT GTG TAA ACT CGA GTT TAC ACA GGA ATT GAG TCG GTT TTT G-3ʹ, shCHAF1A#1 oligo2: 5ʹ-AAT TCA AAA ACC GAC TCA ATT CCT GTG TAA ACT CGA GTT TAC ACA GGA ATT GAG TCG G-3ʹ; shCHAF1A#2 oligo1: 5ʹ-CCG GGA TAC TTG AAC CGA CTC AAT TCT CGA GAA TTG AGT CGG TTC AAG TAT CTT TTT G-3ʹ, shCHAF1A#2 oligo2: 5ʹ-AAT TCA AAA AGA TAC TTG AAC CGA CTC AAT TCT CGA GAA TTG AGT CGG TTC AAG TAT C-3ʹ) from GenePharma (Shanghai, China), then transferred the oligos into a single inducible lentiviral vector (pLKO-TET-Puro, cleaved by A*ge* Ⅰ) to create pLKO-TET-Puro-shCHAF1A#1 and pLKO-TET-Puro-shCHAF1A#2 expression vector. The targeting sequences of shCHAF1A#1 and shCHAF1A#2 were both in the 3’ UTR of CHAF1A mRNA, which was conducive to the recovery of exogenous CHAF1A. We used the pLKO-TET-Puro-scramble vector containing scramble shRNA as a negative control vector. Next, HEK293T cells were co-transfected with the expression vector (pLKO-TET-Puro-scramble, pLKO-TET-Puro-shCHAF1A#1, and pLKO-TET-Puro-shCHAF1A#2) and the packaging vector (psPAX2 and pMD2.G), and the lentiviral supernatant was collected and infected into cancer cells 48 h later. Finally, we successfully generated doxycycline (DOX)-inducible CHAF1A knockdown in the human cancer cell lines A549 and KYSE510. Cells were treated with 1μg/μl DOX for 48 h to knock down endogenous CHAF1A. In some experiments, two pooled shRNAs were used.

### Plasmids, siRNAs, and chemicals

Cells were transfected with plasmids and siRNAs using Lipofectamine 3000 (L3000015, Thermo Fisher Scientific, Waltham, MA, USA) and Lipofectamine RNAiMAX (13778150, Thermo Fisher Scientific), respectively, using standard protocols. The cDNA clones of human CHAF1A wild-type (WT), N terminal (N), middle domain (M), and C terminal (C) and PIP domain-truncated mutations were extracted from HEK293T cells, amplified using PCR, and subcloned into pcDNA3.1-Flag-C and pEGFP-C1 vectors. The pEnCMV-RAD18-HA and pcDNA3.1-RPA2-HA plasmids were purchased from MiaoLing Plasmid Sharing Platform. siRNAs targeting CHAF1A (siCHAF1A#1: 5ʹ-CAG CCA UGG AUU GCA AAG A-3ʹ, siCHAF1A#2: 5ʹ-CAG AAC GAC AAG UUG GCA U-3ʹ, and siCHAF1A#3: 5ʹ-CUC CGC AGA AUA ACU AAG A-3ʹ) and siRNAs targeting RAD18 (siRAD18#1: 5ʹ-GGU AGA CUC UUU GGC ACU U-3ʹ, siRAD18#2: 5ʹ-GCC CGA GGU UAA UGU AGU U-3ʹ, and siRAD18#3: 5ʹ-GCA GUG AUG CUU AUG GUU U-3ʹ) were purchased from GenePharma (Shanghai, China). In some experiments, three pooled siRNAs were used. Hydroxyurea (H8627) was purchased from Sigma-Aldrich (St. Louis, MO, USA). Doxycycline (T1687) was purchased from TOPSCIENCE (Shanghai, China).

### Western blotting

Western blot analysis was performed using standard protocols, as described previously ^22^. Briefly, samples were heated at 95 °C for 5 –15 min in 1 × sodium dodecyl sulfate (SDS) loading buffer, subjected to electrophoresis in 10% SDS-polyacrylamide gels, and transferred to polyvinylidene difluoride (PVDF) membranes. Blocking and antibody dilution were performed using 5% nonfat milk (or bovine serum albumin, BSA) in phosphate-buffered saline (PBS). After incubation with primary antibodies at 4 °C overnight and secondary antibodies at 25 ℃ for 1.5 h, the membranes were washed three times in 1× PBST (1× PBS with 0.1% Tween −20). Images were acquired using a Chemodoc MP system (Bio-Rad, Hercules, CA, USA) with ImageLab software. The following antibodies were used: rabbit polyclonal anti-PCNA (10205-2-AP, Proteintech, Rosemont, USA; 1:5000), mouse monoclonal anti-PCNA (sc-56, Santa Cruz Biotechnology, Dallas, TX, USA; 1:1000), rabbit monoclonal anti-Ub-PCNA (K164) (13439S, Cell Signaling Technology, Danvers, MA, USA; 1:1000), mouse monoclonal anti-GFP-tag (sc-9996,, Santa Cruz Biotechnology; 1:1000), rabbit polyclonal anti-HA-tag (51064-2-AP, Proteintech; 1:8000), mouse monoclonal anti-CHAF1A (sc-133105, Santa Cruz Biotechnology; 1:1000), rabbit polyclonal anti-CHAF1A (17037-1-AP, Proteintech; 1:1000), rabbit polyclonal anti-RAD18 (18333-1-AP, Proteintech; 1:2000), rabbit polyclonal anti-RPA2 (35869S, Cell Signaling Technology; 1:1000), mouse monoclonal anti-Pol η (sc-17770, Santa Cruz Biotechnology; 1:1000), horseradish peroxidase (HRP)- conjugated beta actin monoclonal antibody (HRP-60008, Proteintech; 1:10000). HRP-conjugated monoclonal GAPDH (HRP-60004, Proteintech; 1:20000), and HRP-conjugated monoclonal alpha tubulin (HRP-66031, Proteintech; 1:10000). Secondary antibodies included goat anti-mouse IgG-HRP (sc-2005, Santa Cruz; 1:5000) and goat anti-rabbit IgG-HRP antibodies (sc-2030, Santa Cruz Biotechnology; 1:5000).

### CoIP assays

CoIP was performed as described previously ^23^. To detect interactions between endogenous proteins, Cells were lysed with immunoprecipitation (IP) buffer (50 mM Tris-HCl pH 7.5, 150 mM NaCl, and 0.5% NP-40) supplemented with a protease inhibitor cocktail (HY-K0010, MCE, Shanghai, China). After sonication and centrifugation at 12 000 rpm at 4 °C for 10 min, the protein concentration of the cell lysates was measured using a Pierce BCA Protein Assay Kit (23225, Thermo Fisher Scientific). Equal amounts of total protein (1 mg) were immunoprecipitated by incubation with protein A/G magnetic beads (HY-K0202, MCE) and indicated antibodies at 4 °C overnight with gentle rocking. To detect interactions between PCNA and exogenous GFP-CHAF1A (WT, ΔPIP1, and ΔPIP2), HEK293T cells were transfected with pEGFP-CHAF1A (WT), pEGFP-CHAF1A (ΔPIP1), pEGFP-CHAF1A (ΔPIP2), or pEGFP-vector plasmids, and total protein was collected after 48 h. To confirm the interaction between CHAF1A, RAD18 and RPA2, HEK293T cells were transfected with pEGFP-CHAF1A (WT), pEGFP-CHAF1A (N), pEGFP-CHAF1A (C), pEGFP-CHAF1A (M), or pEGFP-vector plasmids and pEnCMV-RAD18-HA and pcDNA3.1-RPA2-HA, and total protein was collected after 48 h. Equal amounts of total protein (1 mg) were immunoprecipitated by incubation with protein A/G magnetic beads (HY-K0202, MCE) and anti-GFP antibody (sc-9996, Santa Cruz Biotechnology; 1:50) at 4 °C overnight with gentle rocking. The immunoprecipitated proteins were washed three times with cold IP buffer and denatured with 1× SDS sample buffer.

### Cell fractionation

The cell fractionation experiments were performed as described previously ^24^, with slight modifications. Cells in 10 cm dishes were digested using trypsin, collected by centrifugation at 500 × g for 5 min, washed with PBS, and lysed on ice in 600 μL of hypotonic buffer (10 mM Tris-HCl, pH 7.5, 2 mM MgCl_2_, 3 mM CaCl_2_, 320 mM sucrose, 1 mM dithiothreitol (DTT), and 0.3% NP40) supplemented with protease inhibitor cocktail (HY-K0010, MCE). After 10 min, the cells were centrifuged for 5 min at 2800 × *g* and the supernatant containing cytoplasmic proteins was collected. The pellet was washed three times in hypotonic buffer, incubated with 100 μL nuclear extraction buffer (20 mM 2-[4-(2-hydroxyethyl) piperazin-1-yl] ethanesulfonic acid (HEPES), pH 7.7, 1.5 mM MgCl_2_, 420 mM NaCl, 200 mM ethylene diamine tetraacetic acid (EDTA), 25% glycerol, and 1 mM DTT) supplemented with a protease inhibitor cocktail (HY-K0010, MCE) for 30 min at 4 °C, and centrifuged at 8 000 × *g* for 15 min. The supernatant containing soluble nuclear proteins was collected and the chromatin pellet was resuspended in 1× SDS loading buffer, sonicated, and subjected to western blot analysis.

### DNA fiber assays

The DNA fiber assay was performed as described previously ^25^. For monitoring replication fork progression, cells were first incubated with 33 μΜ CldU (C6891, Sigma-Aldrich) for 30 min and then with 330 μM IdU (I7125, Sigma-Aldrich) in the presence of 50 μΜ HU for 30 min. Alternatively, the cells were labeled with CldU for 30 min, exposed to 2 mM HU for 2 h, and then incubated with IdU for 30 min before being harvested in PBS. Labeled cells were quickly trypsinized, resuspended in ice-cold PBS at a density of approximately 1 × 10^6^ cells/mL, and spotted onto a pre-cleaned glass slide (2.5 μL) for lysis in 7.5 μL of lysis buffer (0.5% SDS in 200 mM Tris-HCl (pH 7.4) with 50 mM EDTA). After 5 min, the slides were tilted at 25 °C relative to the horizontal and the resulting DNA spreads were air-dried and fixed in 3:1 methanol/acetic acid for 20 min. After rehydration in PBS, the samples were denatured in 2.5 M HCl for 1 h, washed with PBS, and blocked with 2% BSA in PBS (w/v) containing 0.1% Tween-20 for 40 min. Immunodetection was performed using the following primary antibodies: anti-CldU (rat monoclonal anti-5-bromo-2-deoxyuridine (BrdU)/CldU; BU1/75 ICR1, Abcam, Cambridge, UK; 1:200) and anti-IdU (mouse monoclonal anti-BrdU/IdU; clone B44, BD Biosciences, NJ, USA; 1:25). The samples were then incubated with the following secondary antibodies in a humidified chamber for 1 h at 25 °C: Alexa Fluor® 647 AffiniPure donkey anti-mouse (715-605-150, Jackson ImmunoResearch, West Grove, PA, USA; 1:200) or goat anti-rat Alexa Fluor 488 (A-11006, Invitrogen, San Francisco, CA, USA; 1:200). Images were acquired using a Zeiss (Jena, Germany) LSM 800 fluorescence microscope at 40× magnification and analyzed using ImageJ software (version 1.52). A minimum of 100 individual fibers were analyzed per experiment. Statistical analyses were performed using Prism (version 8.0.2, GraphPad).

### Cell death assays

Cell death was determined by flow cytometry with propidium iodide (P4170, sigma) staining as described previously ^26^. First, cells were seeded onto 12-well plates at a density of 70-80%. The next day, HU was added to the cells and incubated for 48 h. Then, floating and adherent cells were collected and stained with 5 μg/ml propionate iodide at 37 °C for 30 min. The percentage of propionate iodide positive dead cell populations was analyzed using BD Accuri C6 PLUS (BD Biosciences) flow cytometry. At least 8 000 single cells were analyzed per well, and all experiments were at least triplicate.

### PLA assays

PLA assays was performed using Duolink PLA technology (Sigma-Aldrich) according to the manufacturer’s instructions. Briefly, cells were washed once with 1× PBS and treated with cytoskeletal (CSK) extraction buffer [0.2% Triton X-100, 20 mM HEPES-KOH (pH 7.9), 100 mM NaCl, 3 mM MgCl_2_, 300 mM sucrose, 1 mM ethylenebis (oxyethylenenitrilo) tetraacetic acid (EGTA)] containing 4% formaldehyde at 25 °C for 10 min. The cells were then washed three times with 1× PBS, permeabilized with 0.5% NP-40 in 1× PBS for 5 min, and blocked with 5% BSA in PBS at 25 °C for 1 h. After incubation with the indicated primary antibodies at 4 °C overnight, the cells were washed three times with 1× PBS and incubated with anti-mouse and anti-rabbit plus PLA probes (PLA kit, Sigma-Aldrich) at 37 °C for 1 h. The PLA reaction was performed using Duolink in Situ Detection Reagents (PLA kit) according to the manufacturer’s instructions. Finally, the cells were washed three times with buffer B, stained with 4’,6-diamidino-2-phenylindole (DAPI) during the second wash, and mounted with antifade mounting medium (Beyotime) on slides that were sealed with nail polish. Images were captured using a Zeiss LSM 800 fluorescence microscope (40×) and quantified using ImageJ software.

### Colony formation assay

Colony formation assays were performed as described previously ^22^. Briefly, DOX - inducible CHAF1A-knockdown cells were plated at low densities for 48 h in medium containing 1μg/μl DOX, and treated with the indicated doses of HU for 24 h. Then, the cells were replaced with fresh medium and incubated for 7–14 days at 37 °C with 5% CO_2_. After washing with PBS, the cultures were fixed with methanol for 20 min and stained with crystal violet overnight. Colonies were imaged using a Chemodoc MP system (Bio-Rad, Hercules, CA, USA) and analyzed using ImageLab (Bio-Rad, Hercules, CA, USA) and ImageJ software. Each experiment was performed in triplicate.

### Micronuclei analysis

Cells were grown on sterile glass coverslips, treated with HU for 24 h, then fixed in 4% paraformaldehyde/1×PBS for 10 min and permeable in 0.5% Triton X-100/1×PBS for 5 min. The coverslips were then stained with DAPI (Sigma-Aldrich), dehydrated in ethanol, and then sealed. Samples were imaged using Zeiss LSM 800 fluorescence microscope (40×) and scored with the naked eye. Ten photographs were taken in each experiment and the percentage of nuclei with micronuclei in each photograph was calculated separately. Each experiment was performed in triplicate.

### Quantification and statistical analysis

All statistical analyses were performed using Prism (version 8.0.2, GraphPad). Significant differences were determined using *P* values (\**p* < 0.05, \*\**p* < 0.01). Comparisons between two groups were made using unpaired two-tailed *t*-tests. All plotted values represent the mean ± standard deviation (SD). Data in graphs displaying fold changes represent the mean ± SD of fold changes calculated from the mean of control samples.

## Results

### High expression of CHAF1A is associated with poor prognosis in cancer patients

First, we tested the correlation between CHAF1A mRNA levels and overall survival in cancer patients. In a cohort of nearly 2 000 lung cancer patients, high CHAF1A mRNA expression in tumors strongly predicted a poorer overall survival (Figure 1A). Similarly, in a cohort of 364 liver cancer patients, elevated CHAF1A mRNA levels robustly correlated with decreased overall survival (Figure 1B).

**Figure 1.**
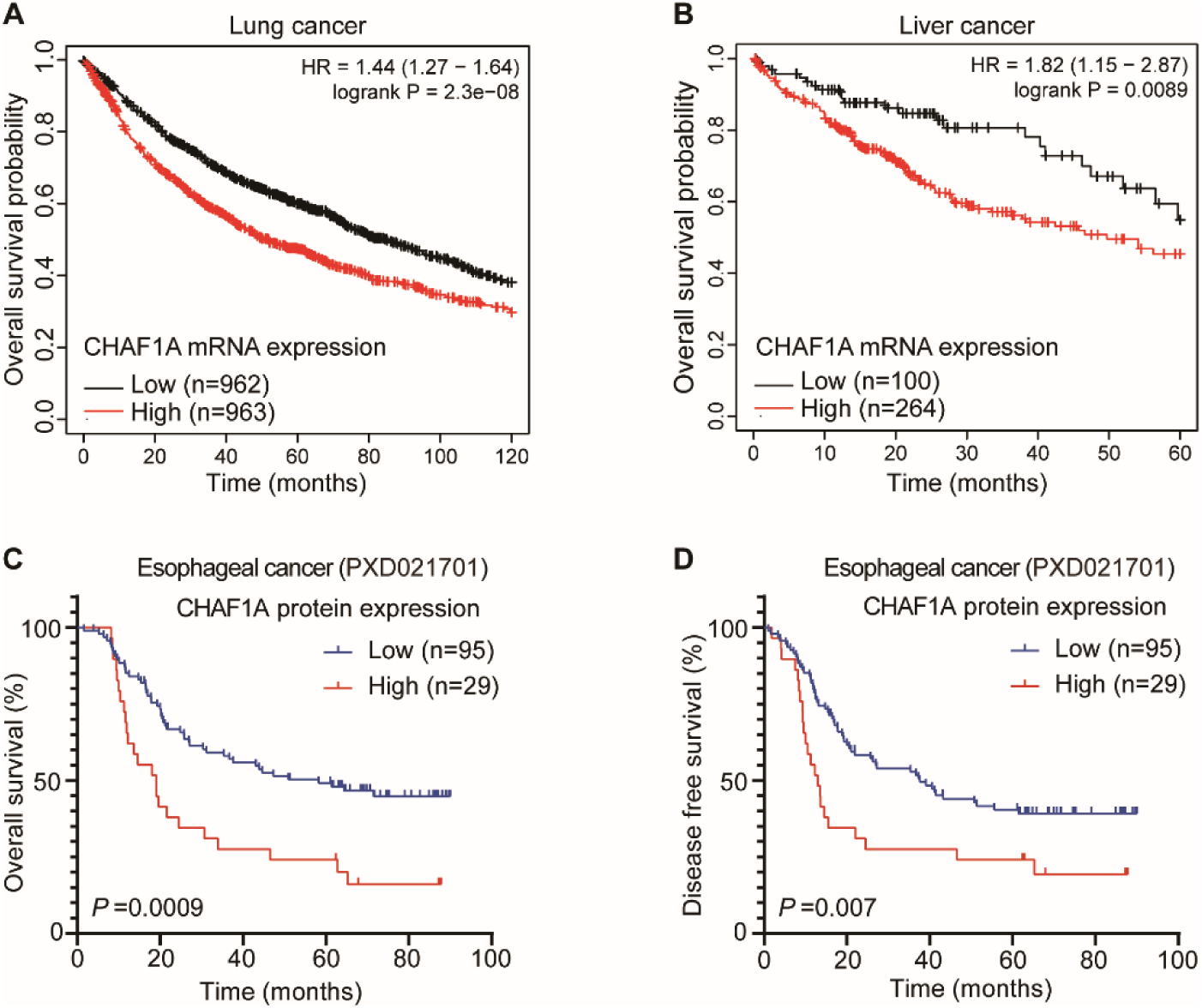
High expression of CHAF1A reduces the survival rate of cancer patients. (A and B) Human lung and liver cancer patients with high CHAF1A mRNA expression levels have worse clinical outcomes. Kaplan-Meier survival was analyzed based on expression levels of CHAF1A mRNA in the cohorts of lung (A) and liver (B) cancer patients using online software (http://kmplot.com/analysis/). Log-rank p values are shown. (C and D) To explore the clinical significance of CHAF1A protein expression cancer, we analyzed the clinicopathological characteristics of 124 patients with esophageal cancer in the esophageal cancer proteome dataset. The effect of CHAF1A protein expression on the overall survival (C) and disease-free survival (D) of patients with EC was evaluated using Kaplan-Meier survival analysis. Since high CHAF1A protein expression are correlated with a poor prognosis in patients with esophageal cancer.

Recently, we performed proteomic analysis on 124 pairs of esophageal cancer tissues and corresponding adjacent non-tumor tissues to identify abnormal changes in protein expression, phosphorylation, and pathways in esophageal cancer ^27^. Next, we examined the association between CHAF1A protein levels and cancer patient survival. Kaplan-Meier survival analysis revealed that esophageal cancer patients with high CHAF1A protein expression had significantly lower overall survival and disease-free survival (Figure 1C and D). Together, these results suggest that cancer patients with high expression of CHAF1A have poor prognosis.

### CHAF1A enhances cancer cell survival under DNA replication stress

CHAF1A possesses important functions in normal DNA replication, but its role under DNA replication stress is unclear. HU is an antitumor drug that blocks DNA replication by inhibiting ribonucleotide reductase ^28^. In order to verify the effect of CHAF1A on the survival ability of cancer cells under DNA replication stress, we treated esophageal cancer cell KYSE510 and lung cancer cell A549 with different doses of HU and measured cell survival with a colony formation assay. We constructed KYSE510 and A549 cell lines with stable lentiviral knockdown of CHAF1A (Figure 2C). The results showed that CHAF1A-knockdown KYSE510 cells and A549 cells were more sensitive to HU compared to control cells (Figure 2A, 2B, and 2D). Next, we used cell death assays to further verify the effect of CHAF1A deficiency on cancer cell survival under DNA replication stress. Cells were treated with HU for 48 h before PI staining, and then the proportion of dead cells was detected by flow cytometry analysis. The results showed that the mortality of CHAF1A-knockdown KYSE510 cells and A549 cells were significantly increased after HU treatment (Figure 2E, 2F, 2G, and 2H). Together, these results demonstrate that CHAF1A enhances the survival of cancer cells under DNA replication stress.

**Figure 2.**
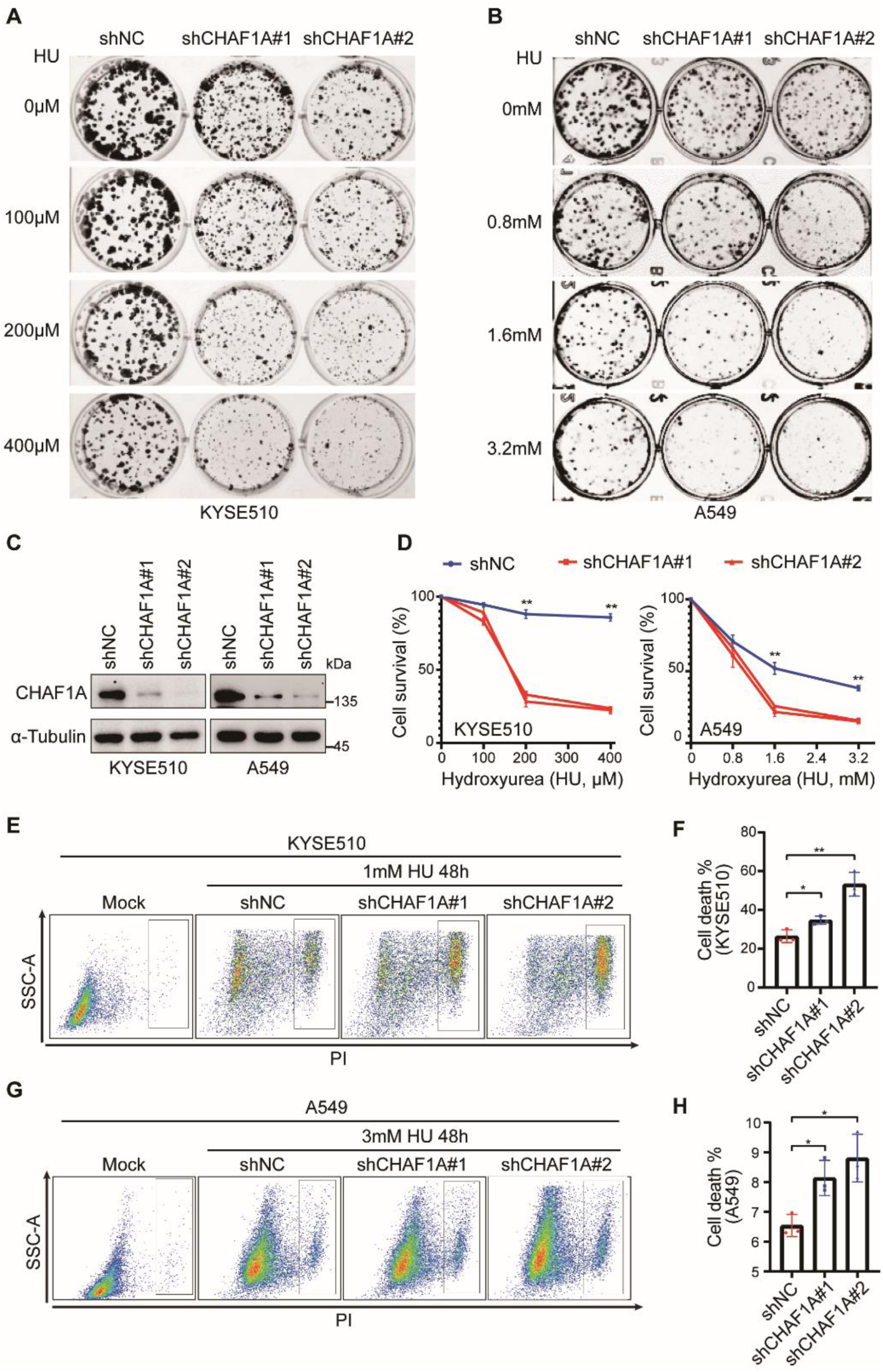
CHAF1A knockdown accelerated HU-induced cancer cell death. (A) Colony formation assay for KYSE510 cells with treatment of the indicated dose of HU. The shNC group was seeded with 500 cells, and the shCHAF1A#1 and shCHAF1A#2 groups were seeded with 1 000 cells respectively. (B) Colony formation assay for A549 cells with treatment of the indicated dose of HU. The shNC group was seeded with 500 cells, and the shCHAF1A#1 and shCHAF1A#2 groups were seeded with 1 000 cells respectively. (C) The CHAF1A knockdown in KYSE510 cells and A549 cells was detected by western blotting. (D) Statistical plots of colony formation assay in (A) and (B) (n = 3). (E) Cell death measurements in KYSE510 cells treated with 1mM HU for 48h. Data are plotted in (F) (n = 3). (G) Cell death measurements in A549 cells treated with 3mM HU for 48h. Data are plotted in (H) (n = 3).

### CHAF1A facilitates the restart of stalled replication forks and reduces genomic instability

The above results suggest that CHAF1A enhances cancer cells’ tolerance to DNA replication stress, so we proposed that CHAF1A might further protect the replication forks and genomic stability under DNA replication stress. We first used DNA fiber assays to directly examine the impact of CHAF1A on DNA replication forks under replication stress. Nascent DNA was sequentially labeled with the thymidine analogs 5-chloro-2-deoxyuridine (CldU) and 5-iodo-2-deoxyuridine (IdU) with low concentrations of HU (50μM). The ratio of IdU- to CldU-labeled replication tracts were lower in CHAF1A-knockdown cells than in control cells (Figure 3A and 3B). This result indicated that CHAF1A knockdown inhibited DNA replication rate under DNA replication stress.

**Figure 3.**
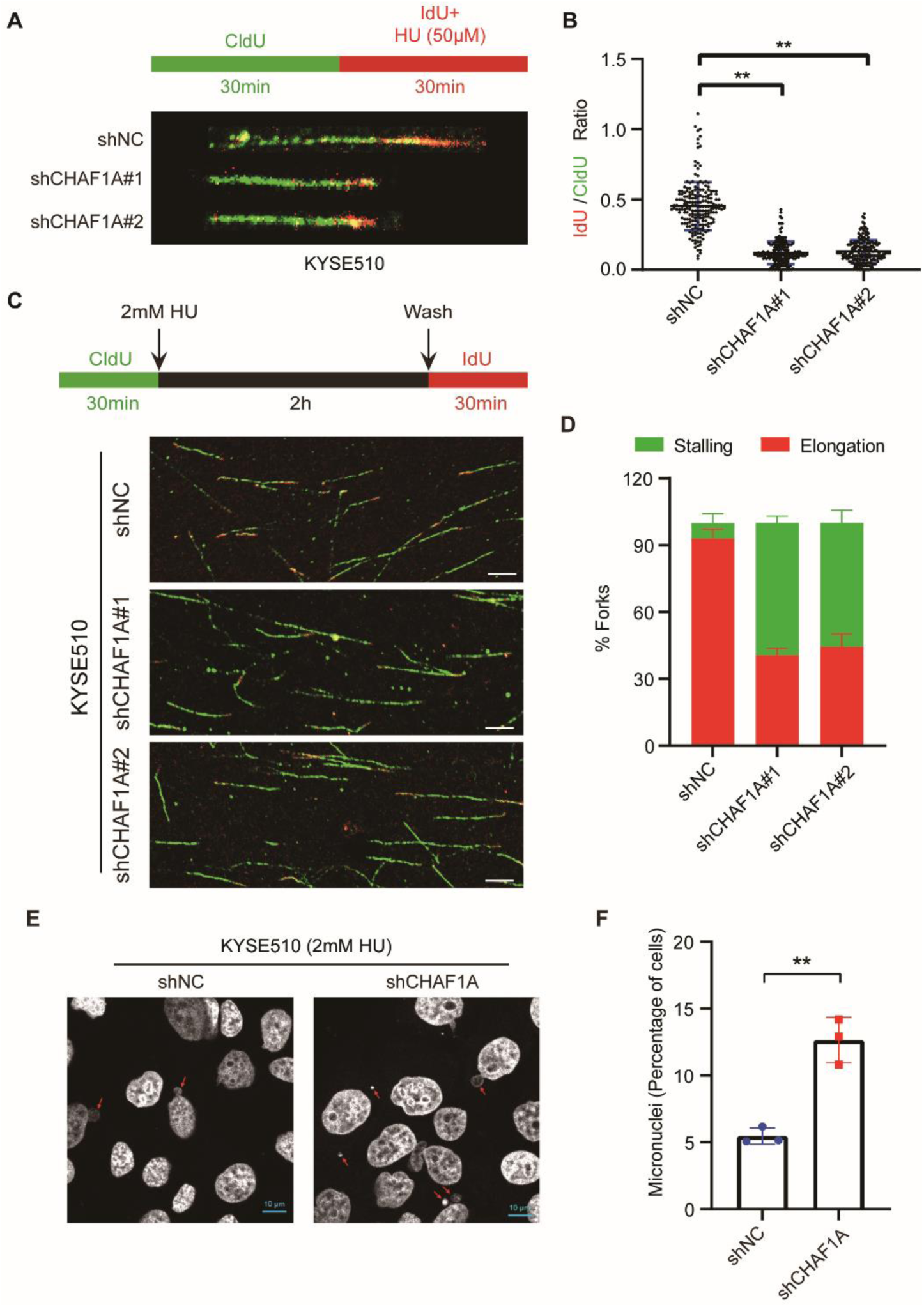
CHAF1A knockdown inhibits stalled replication fork restarting and genome integrity under DNA replication stress. (A) DNA fiber assay was used to measure DNA replication in inducible CHAF1A knockdown KYSE510 cells. Schematic of alternative CldU/IdU pulse-labeling protocol to evaluate fork status. Representative images of DNA fiber assays are shown. Data are plotted in (B) (n = 200). (C) Schematic of alternative CldU/IdU pulse-labeling protocol to evaluate fork status. Representative images of DNA fiber assays are shown. (D) Image J was used to calculate the percentage of stalling forks and elongation forks in each field of view (n = 10). (E) Percentage of cells presenting micronuclei in inducible CHAF1A-knockdown KYSE510 cells (mix shCHAF1A#1 and shCHAF1A#2) treated with 2mM HU for 24h. Data are plotted in (F) (n = 3).

The deceleration of replication forks in CHAF1A-knockdown cells prompted us to ask whether CHAF1A affected the restart of stalled replication forks, which occurs throughout the genome in response to DNA replication stress. To investigate whether CHAF1A knockdown affects replication fork restarting after DNA replication stress, we also performed DNA fiber assay. As reported previously ^29^, HU treatment after CldU labeling reduced IdU detection due to failed replication fork restarting; however, CHAF1A knockdown further reduced replication fork restarting (Figure 3C and 3D), suggesting that CHAF1A facilitates the restarting of stalled replication forks under DNA replication stress.

The restarting of replication forks following DNA replication stress is critical for genomic stability, and continued replication fork stalling may lead to fork collapse and cell death ^16^. To understand the effect of CHAF1A knockdown on genomic instability, we examined the number of micronuclei in cancer cells under replication stress. We observed that CHAF1A-knockdown cancer cells formed more micronuclei than control cells under DNA replication stress (Figure 3E and 3F). Altogether, the results indicate that CHAF1A inhibits genomic instability by promoting DNA replication fork restart under DNA replication stress.

### CHAF1A promotes the TLS pathway induced by HU

TLS pathway protects cancer cells from cell death caused by DNA replication stress inducers, which is one of the important reasons for clinical chemoradiotherapy resistance ^30^. In addition to UV, HU can also significantly activate the TLS pathway ^31, 32^. Therefore, we speculated that CHAF1A may influence HU-induced monoubiquitination of PCNA at K164, which is a key step in the TLS pathway. To test the conjecture, we examined the effects of CHAF1A deficiency on HU-induced PCNA monoubiquitination. Western blotting analysis showed that CHAF1A deficiency significantly inhibited PCNA monoubiquitination induced by HU (Figure 4A and 4B).

**Figure 4.**
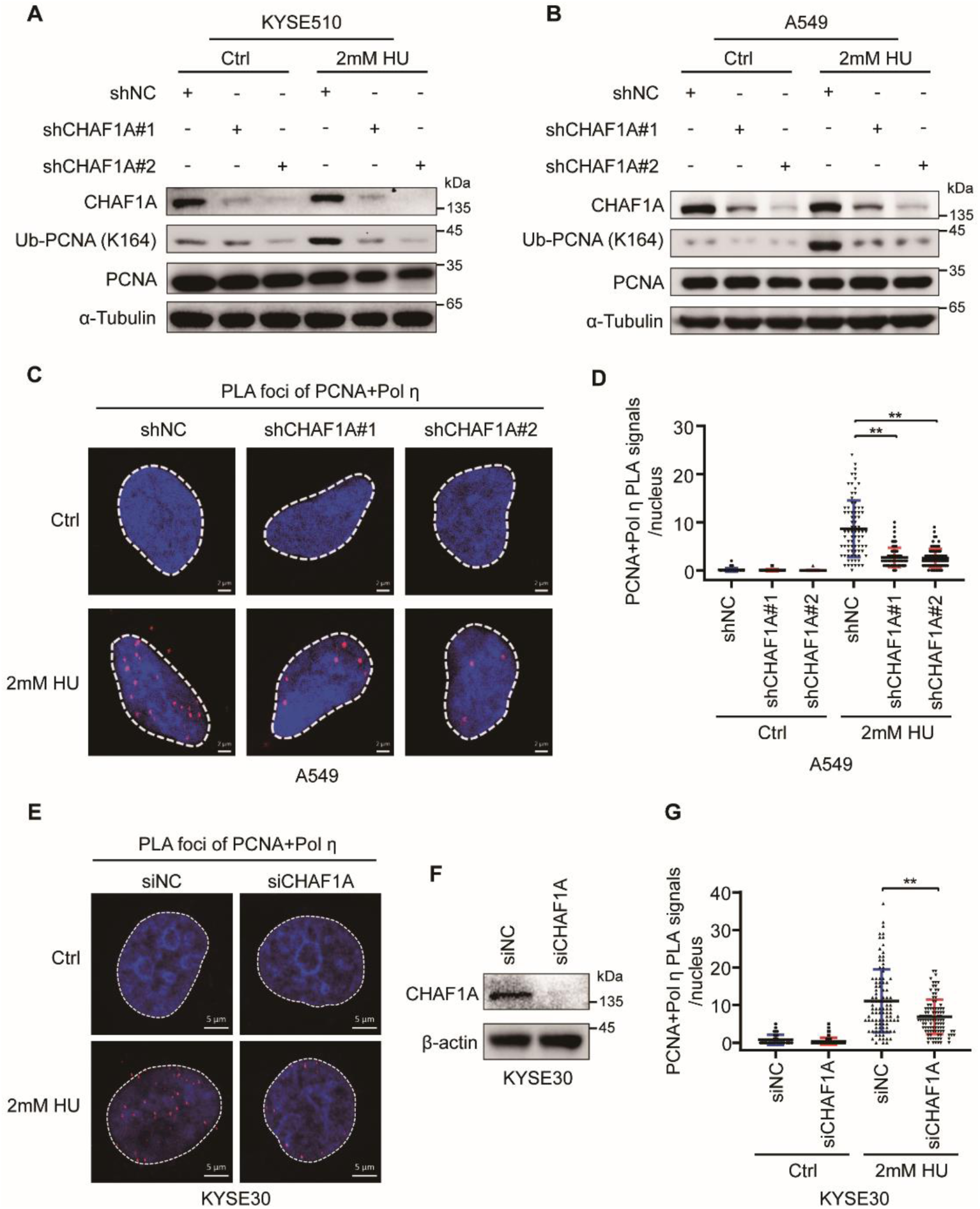
CHAF1A knockdown impedes HU-induced PCNA monoubiquitination and PCNA-Pol η interaction. (A) CHAF1A knockdown KYSE510 cells were treated with or without 2mM HU for 4 h. Whole-cell lysates were analyzed using western blotting with the indicated antibodies. (B) CHAF1A knockdown A549 cells were treated with or without 2mM HU for 4 h. Whole-cell lysates were analyzed using western blotting with the indicated antibodies. (C) CHAF1A knockdown A549 cells were treated with or without 2mM HU for 4 h. PLA assays was used to detect the interaction between PCNA and Pol η (scale bar, 2 μm), and (D) each spot represents the number of PLA foci in an individual nucleus (n = 80). (E) KYSE30 cells were transfected with control or CHAF1A siRNA oligos for 72 h and then treated with or without 2mM HU for 4 h. PLA assays was used to detect the interaction between PCNA and Pol η (scale bar, 5 μm), (F) the CHAF1A knockdown in KYSE30 cells was detected by western blotting, and (G) each spot represents the number of PLA foci in an individual nucleus (n = 100).

Since CHAF1A positively regulated PCNA monoubiquitination, we speculated that CHAF1A may also influence the TLS pathway. To test this hypothesis, we used PLA assays to verify the effect of CHAF1A on the interaction between PCNA and Pol η, which is the best-characterized TLS polymerase in mammalian cells ^33^. Consistently, the interaction between endogenous PCNA and Pol η was only detectable after HU treatment and was reduced in CHAF1A-knockdown cells (Figure 4C and 4D). Moreover, we further demonstrated that CHAF1A is essential for HU-induced Pol η recruitment to PCNA by using siRNA knockdown in esophageal cancer cells KYSE30 (Figure 4E, 4F, and 4G). These results strongly suggest that CHAF1A promotes the HU-induced TLS pathway in response to DNA replication stress.

### CHAF1A promotes PCNA monoubiquitination independent of CHAF1A-PCNA interaction

The interaction between CHAF1A and PCNA promotes histone remodeling during DNA replication ^34^. We wanted to explore whether the effect of CHAF1A on PCNA monoubiquitination was dependent on the interaction between CHAF1A and PCNA. Previous studies have reported that CHAF1A contains two PIP (PCNA-interaction peptide) domains (PIP1 and PIP2) (Figure 5A) ^35^. However, the domain on which CHAF1A interacts with PCNA is still controversial. Ben-Shahar et al. found that CHAF1A interacts with PCNA through its PIP1 domain ^35^, while Cheng et al. demonstrated that CHAF1A interacts with PCNA dependent on the PIP2 domain ^36^. To confirm which PIP domain of CHAF1A affected the interaction with PCNA, we performed CoIP assays with anti-GFP antibody in HEK293T cells transfected with GFP-vector, GFP-CHAF1A (WT), GFP-CHAF1A (ΔPIP1), or GFP-CHAF1A (ΔPIP2). Western blotting analysis showed that CHAF1A interacts with PCNA dependent on the PIP2 domain (Figure 5B). To test this result, we then performed in situ proximity ligation assays (PLA), which are widely used to detect protein interactions in cells with high specificity and sensitivity ^37^. We introduced GFP-vector, GFP-CHAF1A (WT), GFP-CHAF1A (ΔPIP1), or GFP-CHAF1A (ΔPIP2) into KYSE510 cells and performed PLA assay. In contrast to GFP-CHAF1A (WT) and GFP-CHAF1A (ΔPIP1), the GFP-CHAF1A (ΔPIP2) failed to interact with PCNA (Figure 5C and 5D).

**Figure 5.**
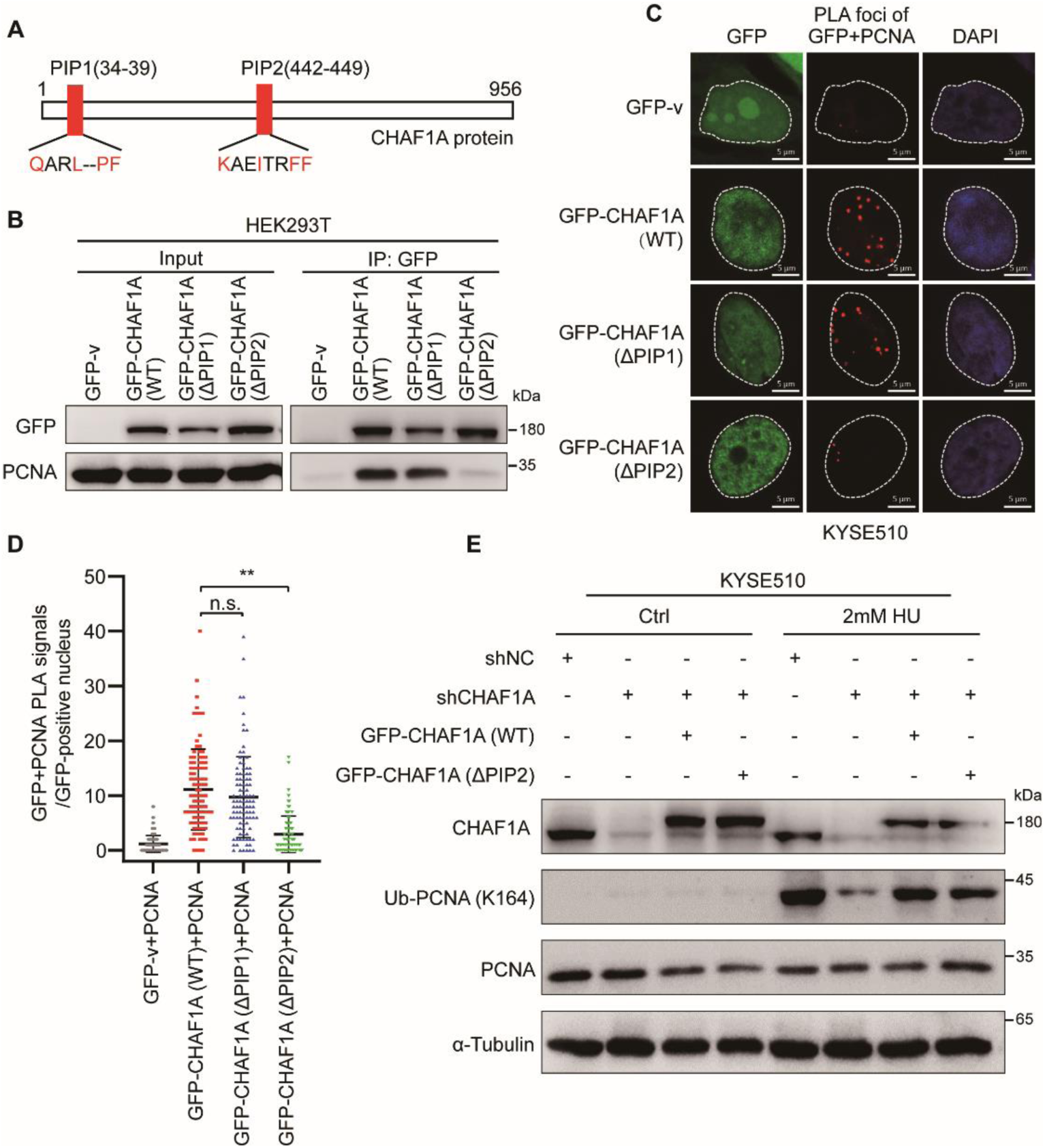
The CHAF1A PIP2 domain mediates the CAHF1A-PCNA interaction, but is not necessary for PCNA monoubiquitination. (A) Schematics of CHAF1A PIP domains. (B) Interactions of GFP-CHAF1A (WT, ΔPIP1, and ΔPIP2) with PCNA in HEK293T cells assessed by CoIP using anti-GFP antibody. ΔPIP1: PIP1 domain deletion, ΔPIP2: PIP2 domain deletion. (C) The in-situ interaction of GFP-CHAF1A (WT, ΔPIP1, and ΔPIP2) with PCNA in KYSE510 cells was evaluated by PLA assay using GFP and PCNA antibodies. GFP-v was used as a negative control (scale bar, 5 μm). (D) The number of PLA foci in GFP-positive nuclei was counted, each spot represents the number of PLA foci in an individual nucleus (n = 100). (E) CHAF1A-knockdown KYSE510 cells (mix shCHAF1A#1 and shCHAF1A#2) were transfected with GFP-tagged CHAF1A (WT and ΔPIP2) vectors and then treated with or without 2mM HU for 4 h. Whole-cell lysates were analyzed using western blotting with the indicated antibodies.

Furthermore, to test the effect of CHAF1A-ΔPIP2 mutant on HU-induced PCNA monoubiquitination, we treated KYSE510 cells with HU to activate PCNA monoubiquitination. Western blotting analysis showed that knockdown of CHAF1A significantly reduced PCNA monoubiquitination, while overexpression of GFP-CHAF1A (WT) and GFP-CHAF1A (ΔPIP2) significantly restored PCNA monoubiquitination (Figure 5E). Together, these results suggest that the interaction of CHAF1A with PCNA requires the PIP2 domain. Meanwhile, the enhancement of PCNA monoubiquitination by CHAF1A was independent of the interaction between CHAF1A and PCNA.

### CHAF1A stimulates PCNA monoubiquitination in a RAD18-dependent manner

Next, we wanted to elucidate the molecular mechanism by which CHAF1A promotes HU-induced PCNA monoubiquitination. RAD18 is an E3 ubiquitin ligase that directly mediates the monoubiquitination of PCNA at K164 during TLS ^38^. Here, we hypothesized that CHAF1A affected RAD18-mediated PCNA monoubiquitination. To test this hypothesis, KYSE510 cells transfected with GFP-CHAF1A (WT) were treated with HU, and western blotting analysis showed that CHAF1A overexpression significantly increased PCNA monoubiquitination, which was inhibited by RAD18 knockdown (Figure 6A). This result indicated that CHAF1A-mediated PCNA monoubiquitination is dependent on RAD18.

**Figure 6.**
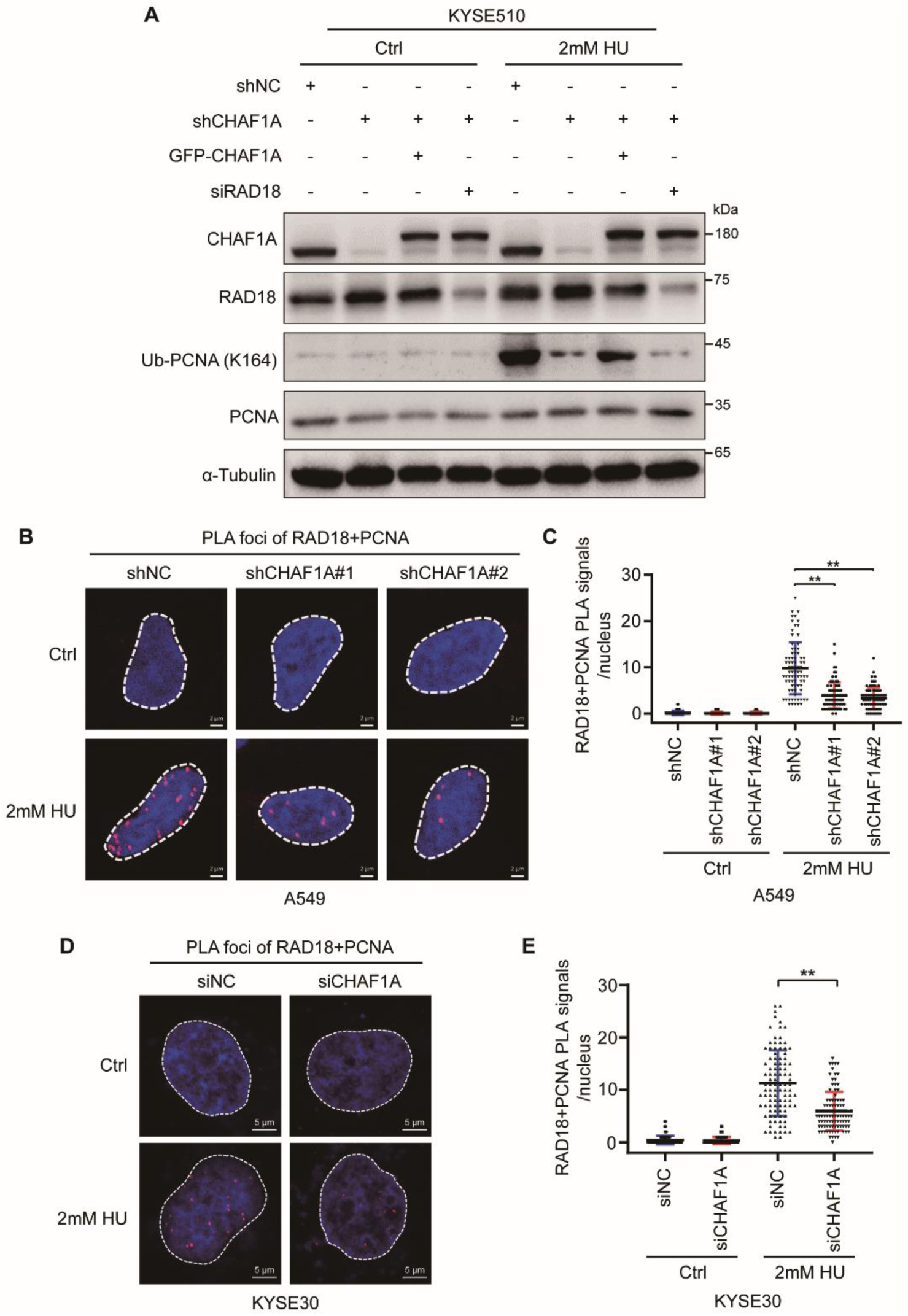
CHAF1A promotes the interaction between RAD18 and PCNA in response to DNA replication stress. (A) CHAF1A-knockdown KYSE510 cells (mix shCHAF1A#1 and shCHAF1A#2) were transfected with indicated siRNAs before transfection with plasmid encoding GFP-CHAF1A and then treated with or without 2mM HU for 4 h. (B) CHAF1A-knockdown A549 cells were treated with or without 2mM HU for 4 h. PLA assay was used to detect the interaction between RAD18 and PCNA (scale bar, 2 μm), and (C) each spot represents the number of PLA foci in an individual nucleus (n = 80). (D) KYSE30 cells were transfected with control or CHAF1A siRNA oligos for 72 h and then treated with or without 2mM HU for 4 h. PLA assays was used to detect the interaction between RAD18 and PCNA (scale bar, 5 μm), (E) The number of PLA foci was counted; each spot represents the number of PLA signals in an individual nucleus (n = 100).

Next, we hypothesized that CHAF1A affected the interaction between RAD18 and PCNA, which is a necessary step for RAD18 to monoubiquitinate PCNA. To verify this conjecture, we performed PLA assays to detect the effect of CHAF1A on the interaction between PCNA and RAD18. We found that the interaction between PCNA and RAD18 was significantly enhanced by HU treatment, and was reduced in CHAF1A-knockdown A549 cells (Figure 6B and 6C). In addition, using siRNA knockdown in esophageal cancer cells KYSE30, we further demonstrated that CHAF1A is essential for HU-induced RAD18 recruitment to PCNA (Figure 6D and 6E). Together, these results demonstrate that CHAF1A promotes the PCNA monoubiquitination by recruiting RAD18 to PCNA.

### CHAF1A mediated the interaction between RAD18 and RPA2 under DNA replication stress

Next, we wanted to know how CHAF1A facilitated RAD18 recruitment to PCNA. The major ssDNA-binding protein RPA interacts with RAD18 to mediate the PCNA monoubiquitination ^39^. Interestingly, sequence alignments identified an evolutionarily conserved motif within CHAF1A protein consisting of amino acids 409-418 (KxRQxALxxK) that closely resembles the RPA2-binding motif of other RPA2-interacting proteins including ETAA1, SMARCAL1, TIPIN and XPA ^40–42^ (Figure 7A). We hypothesize that CHAF1A is an RPA2 interacting protein that mediates RAD18 interaction with RPA. To test this hypothesis, we first performed CoIP using anti-CHAF1A antibody in KYSE510 cells and A549 cells, and found that CHAF1A interacts with RAD18 and RPA2 (Figure 7B and 7C). Subsequently, the interaction of the CHAF1A with RAD18 and RPA2 was further validated by PLA assay (Figure 7D and 7E). To further confirm the interaction domain between CHAF1A, RAD18 and RPA2, we constructed GFP-tagged expression vectors of the N-terminal (N), middle domain (M), and C-terminal (C) of CHAF1A protein, respectively (Figure 7F) and performed CoIP assays using anti-GFP antibody in HEK293T cells. We found that the middle domain of CHAF1A mediates binding to RPA2, whereas the C-terminal domain of CHAF1A mediates binding to RAD18 (Figure 7G).

**Figure 7.**
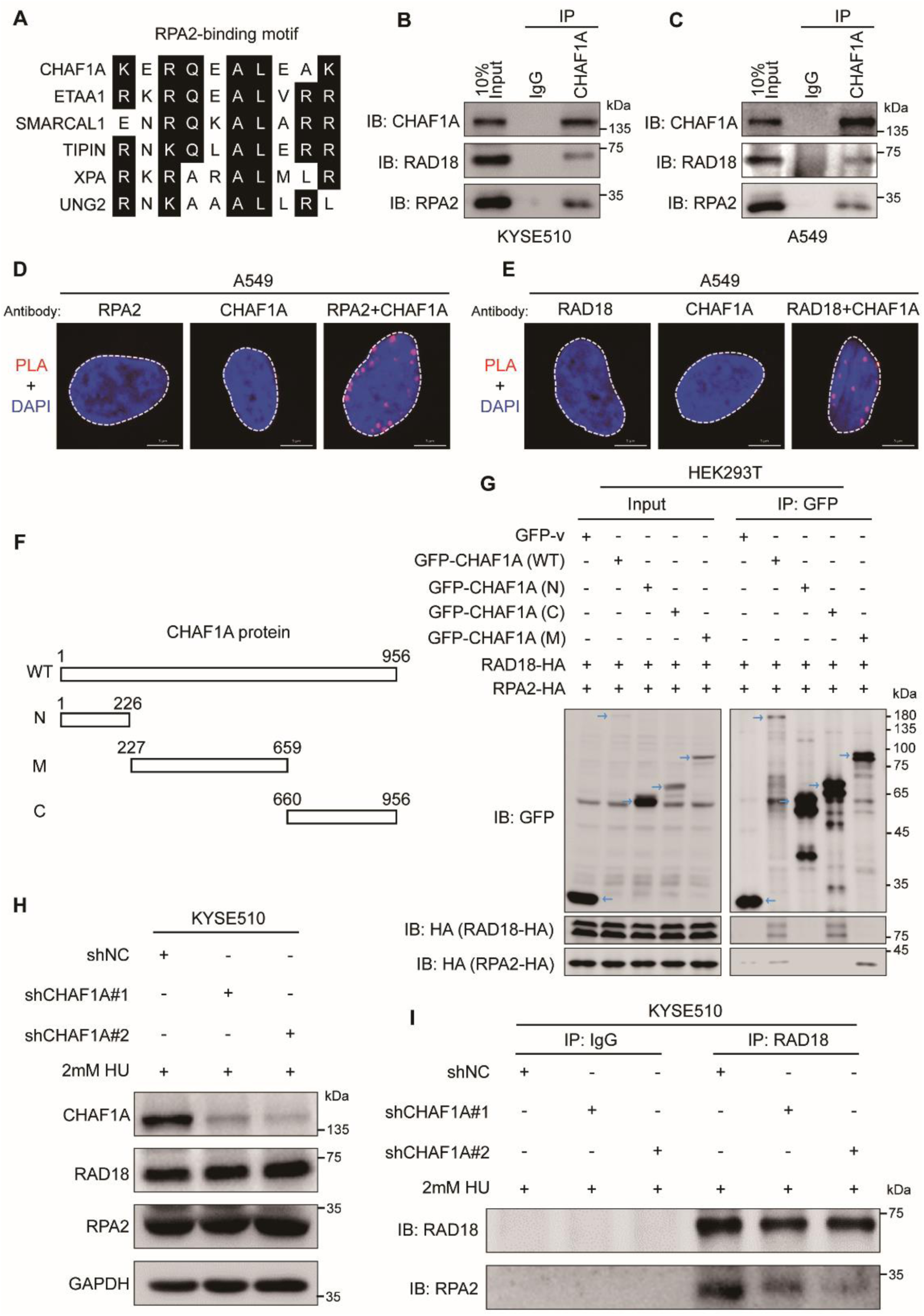
CHAF1A, RPA2 and RAD18 interact with each other in response to DNA replication stress. (A) Sequence alignment of CHAF1A 409-418 with the RPA2-binding motif of other RPA2-interacting proteins. (B) Association of endogenous CHAF1A with RPA2 and RAD18 in KYSE510 cells assessed by CoIP using anti-CHAF1A antibodies. (C) Association of endogenous CHAF1A with RPA2 and RAD18 in A549 cells assessed by CoIP using anti-CHAF1A antibodies. The input sample is diluted to 10%. (D) PLA assay for the interaction of CHAF1A and RPA2 in A549 cells (scale bar, 5 μm). (E) PLA assay for the interaction of CHAF1A and RAD18 in A549 cells (scale bar, 5 μm). (F) Schematic diagram of the CHAF1A mutants. WT, wild type. N, N terminal. M, middle domain. C, C terminal. (G) Interactions of GFP-CHAF1A (WT, N, C, and M) with RAD18-HA and RPA2-HA in HEK293T cells assessed by CoIP using anti-GFP antibodies. The blue arrow indicates the target protein band. (H) CHAF1A knockdown KYSE510 cells were treated with 2mM HU for 4 h. Whole-cell lysates were analyzed using western blotting with the indicated antibodies. (I) CHAF1A knockdown KYSE510 cells were treated with 2mM HU for 4 h, then endogenous RAD18 with RPA2 assessed by CoIP using anti-RAD18 antibodies.

Since the interaction between RAD18 and RPA2 can regulate PCNA monoubiquitination ^39^, we conjectured that CHAF1A mediates the interaction between RAD18 and RPA2 under DNA replication stress. After HU treatment, the protein levels of RAD18 and RPA2 were not affected by the knockdown of CHAF1A (Figure 7H). Interestingly, the HU-induced interaction between RAD18 and RPA2 was significantly inhibited by CHAF1A knockdown (Figure 7I). Together, CHAF1A is essential for the interaction of RAD18 with RPA2 under DNA replication stress.

## Discussion

Endogenous and exogenous DNA replication stress can damage rapidly proliferating cancer cells; however, these cells can bypass DNA damage to maintain genomic stability and survive via the TLS pathway mediated by PCNA monoubiquitination ^8^. CHAF1A traditionally functions as a PCNA partner for chromatin remodeling functions and is involved in histone assembly ^17^. In our study, we identified a new function of CHAF1A that plays an important regulatory role in the TLS pathway. Specifically, DNA replication stress causes replication fork stalling. CHAF1A interacts with RPA2 and RAD18 to promote the recruitment of RAD18 to PCNA, thus enhancing the monoubiquitination of PCNA. Once PCNA is monoubiquitinated, the Y-family DNA polymerase Pol η replaces Pol δ/ε to allow stalled replication forks to proceed. High expression of CHAF1A enables cancer cells to enhance the TLS pathway in response to endogenous and exogenous DNA replication stress, improve the survival ability of cancer cells, and then promote cancer progression (Figure 8).

**Figure 8.**
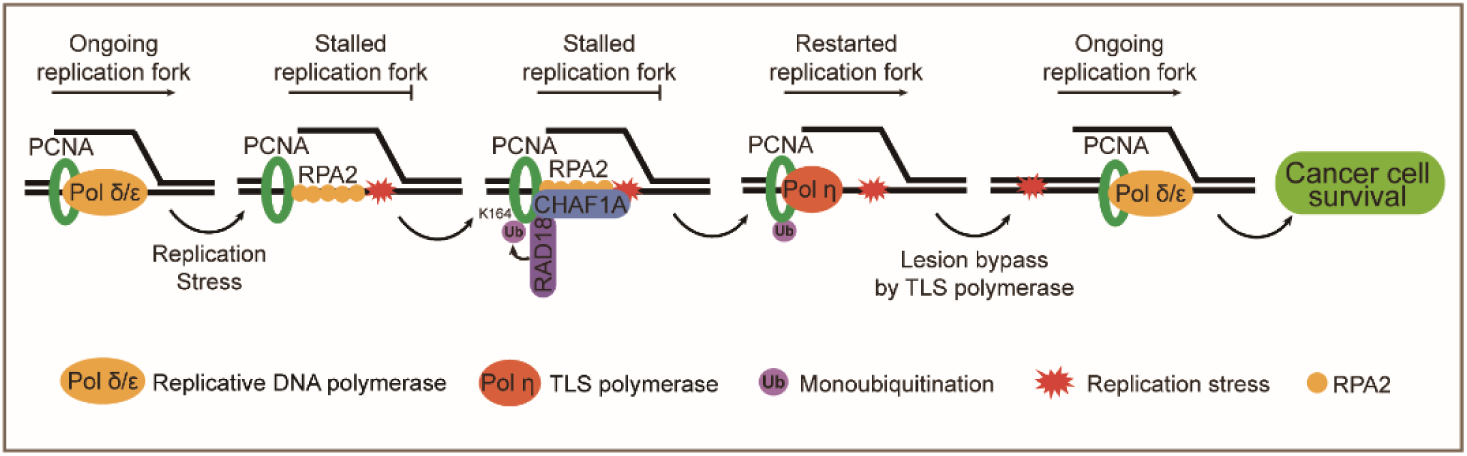
Model depicting the molecular function of CHAF1A in regulating TLS.

CHAF1A is a subunit of the CAF-1 complex, which is recruited to the DNA replication and DNA damage sites by PCNA and is responsible for histone assembly and chromatin remodeling ^43^. During DNA replication stress, PCNA monoubiquitination rapidly initiates the TLS pathway, which recruits lesion-tolerant Y-family polymerases to stalled replication sites ^7^. Dungrawala et al. found that most PCNA and CAF-1 molecules were dissociated from stalled forks under replication stress, but PCNA and CAF-1 were present on stalled forks during a long period of DNA replication stress ^44^. These results suggest that PCNA and CAF-1 complex may respond synergistically to replication stress, but the specific molecular mechanism remains unclear. Moreover, previous studies revealed that CHAF1A contains two PIP domains by sequence alignment, which may mediate the interaction between CHAF1A and PCNA ^35^. Nevertheless, previous studies have shown that CHAF1A interacts with PCNA depending on which PIP domain is controversial ^36^. Therefore, in this study, we demonstrated that CHAF1A interacts with PCNA through the PIP2 domain. Furthermore, we found that CHAF1A promotes the TLS pathway in response to DNA replication stress by regulating PCNA monoubiquitination. Unexpectedly, our data showed that CHAF1A promotes PCNA monoubiquitination independent of the interaction between CHAF1A and PCNA. These results suggest that the involvement of CHAF1A in PCNA monoubiquitination is independent of its traditional histone assembly function.

RAD6/RAD18 is a pair of E2/E3 ubiquitin ligases that mediate monoubiquitination of PCNA at K164 ^38^. Although there are approximately 10 E2 enzymes and more than 60 E3 ligases in eukaryotes, RAD6/RAD18 is the major E2/E3 pair responsible for PCNA monoubiquitination during TLS ^38^. In our study, we demonstrated that CHAF1A promotes PCNA monoubiquitination dependent on RAD18. Moreover, CHAF1A enhances the binding of RAD18 to PCNA. RPA is a heterotrimeric complex composed of RPA1 (also known as RPA70), RPA2 (also known as RPA32), and RPA3 (also known as RPA14), and binds to ssDNA with a very high affinity ^45^. Previous studies have reported that recruitment of RAD6/RAD18 complexes at stalled replication forks is RPA-dependent ^39^. Thus, CHAF1A may promote RAD18 binding on chromatin in a PCNA-independent manner. Here, we found that CHAF1A, RAD18 and RPA2 interact with each other. Under DNA replication stress, CHAF1A promotes the binding of RAD18 on chromatin by mediating the interaction between RAD18 and RPA2, thereby promoting PCNA monoubiquitination.

Prolonged DNA replication stress is thought to lead to replication fork collapse ^46^. Importantly, our study demonstrates that CHAF1A plays a crucial role in fork restarting under DNA replication stress. Under DNA replication stress, CHAF1A-knockdown cells had accumulated more stalled replication forks than control cells, suggesting that CHAF1A is essential for maintaining replication fork integrity and genome stability in cancer cells under DNA replication stress. Cancer cells exhibit strong DNA replication stress and DNA repair responses that can make them resistant to chemotherapeutic drugs targeting DNA replication. Therefore, inhibiting the DNA damage and DNA replication stress responses has become an attractive therapeutic strategy enhancing chemotherapy sensitivity in patients with drug-resistant cancers ^30^. Moreover, high expression of CHAF1A is closely associated with poor prognosis in cancer patients, indicating that high expression of CHAF1A promotes cancer progression. Consistently, we found that inhibiting CHAF1A enhanced the sensitivity of cancer cells to HU, suggesting that CHAF1A is a potential target for cancer therapy.

In summary, our study identified CHAF1A as a key regulator of TLS pathway in cancer cells and CHAF1A inhibits genomic instability by regulating the TLS pathway, promoting cancer cell viability and cancer progression. CHAF1A promotes fork restart under DNA replication stress. Mechanistically, we demonstrated that CHAF1A promotes PCNA monoubiquitination in a RAD18-dependent manner, and this process is independent of CHAF1A-PCNA interaction. Moreover, we revealed that CHAF1A interacts with RAD18 and RPA2, mediating the chromatin binding of RAD18 under DNA replication stress. Together, the findings of this study enrich our understanding of the regulatory mechanisms underlying the TLS pathway and provide insights into the relationship between CHAF1A and the malignant progression of cancer.

## Acknowledgments

This work was supported by the National Natural Science Foundation of China (82273108 and 82173034), the Innovative Team Grant of Guangdong Department of Education (2021KCXTD005), the Science and Technology Special Fund project of Guangdong Province in 2021 (No. 210729156901797) and the 2020 Li Ka Shing Foundation Cross-Disciplinary Research Grant (2020LKSFG07B).

## Data availability

The lung cancer and liver cancer’s clinical data analyzed in this study were obtained from online software at http://kmplot.com/analysis/. The esophageal cancer proteome data were obtained from a previous paper (doi: 10.1038/s41467-021-25202-5). All source data are provided in this study.

## Author contributions

Bing Wen: conceptualization, methodology, investigation, data curation, and writing-original draft. Hai-Xiang Zheng: formal analysis and investigation. Dan-Xia Deng: software and visualization. Zhi-Da Zhang: formal analysis and visualization. Jing-Hua Heng: software and formal analysis. Lian-Di Liao: resources. Li-Yan Xu: funding acquisition, writing-review and editing. En-Min Li: project administration and supervision.

